# Solanaceae specialized metabolism in a non-model plant: trichome acylinositol biosynthesis

**DOI:** 10.1101/2020.03.04.977181

**Authors:** Bryan J. Leong, Steven M. Hurney, Paul D. Fiesel, Gaurav D. Moghe, A. Daniel Jones, Robert L. Last

**Affiliations:** Department of Plant Biology, Michigan State University, East Lansing, MI, USA; Department of Chemistry, Michigan State University, East Lansing, MI, USA; Department of Biochemistry and Molecular Biology, Michigan State University, East Lansing, MI, USA

## Abstract

Plants make hundreds of thousands of biologically active specialized metabolites varying widely in structure, biosynthesis and the processes that they influence. An increasing number of these compounds are documented to protect plants from harmful insects, pathogens, or herbivores, or mediate interactions with beneficial organisms including pollinators and nitrogen fixing microbes. Acylsugars – one class of protective compounds – are made in glandular trichomes of plants across the Solanaceae family. While most described acylsugars are acylsucroses, published examples also include acylsugars with hexose cores. The South American fruit crop *Solanum quitoense* (Naranjilla) produces acylsugars that contain a *myo*-inositol core. We identified an enzyme that acetylates triacylinositols, a function homologous to the last step in the *Solanum lycopersicum* acylsucrose biosynthetic pathway. Our analysis reveals parallels between *S. lycopersicum* acylsucrose and *S. quitoense* acylinositol biosynthesis, suggesting a common evolutionary origin.

**Material availability:** The author responsible for distribution of materials integral to the findings presented in this article in accordance with the policy described in the Instructions for Authors (www.plantphysiol.org) is: Robert L. Last.

**One sentence summary:** Evidence that the final step in *Solanum quitoense* acylinositol biosynthesis evolved from an acylsucrose acetyltransferase enzyme.

## Introduction

Plants are master chemists and collectively produce an array of structurally diverse specialized metabolites (traditionally called secondary metabolites) from common building blocks (Bennett and Wallsgrove, 1994; Verpoorte and Alfermann, 2000; Wink, 2010). Examples include alkaloids derived from amino acids (Ziegler and Facchini, 2008), terpenes made from isoprenoids (Chappell, 1995; Pichersky and Raguso, 2018; Zhou and Pichersky, 2020) and acylsugars synthesized from acyl-CoAs and sugars (Fan et al., 2019). Most specialized metabolites are restricted in cell- or tissue-specific accumulation and found in phylogenetically restricted groups of plants (Pichersky and Lewinsohn, 2011). An increasing number have been shown to be beneficial – examples include deterring or killing microbes and herbivores, inhibiting germination or growth of competitor plants, or attracting beneficial insects and microbes (Bennett and Wallsgrove, 1994; Pichersky and Gershenzon, 2002; Howe and Jander, 2008; Leckie et al., 2016; Massalha et al., 2017). Humans adapted specialized metabolites as medicines including the analgesic morphine and anti-cancer drugs paclitaxel and vinblastine (Fabricant and Farnsworth, 2001), for cooking (for examples, piperine in black pepper and capsaicin in ‘hot’ chilies) (Srinivasan, 2013) and fragrances (e.g. limonene and linalool) (Dudareva and Pichersky, 2006). As biologically active molecules, specialized metabolites can have adverse effects on the plant making them and production or storage of these compounds in specialized structures can mitigate negative effects (Schilmiller et al., 2012b; Schenck and Last, 2019). Epidermal glandular trichomes are an example of biochemical factories that store a wide variety of specialized metabolites across the Plantae kingdom (Fahn, 2000; Schuurink and Tissier, 2019). Classes of specialized metabolites produced in these glandular trichomes include terpenoids (Sallaud et al., 2009; Schilmiller et al., 2009; Brückner et al., 2014), flavonoids (Tattini et al., 2000; Schmidt et al., 2011; Kim et al., 2014), and acylsugars (Schilmiller et al., 2010; Schilmiller et al., 2012a; Schilmiller et al., 2015; Fan et al., 2016).

Acylsugars are specialized metabolites made from sugars and acyl-CoAs in species across the Solanaceae family (King et al., 1986; King and Calhoun, 1988; Matsuzaki et al., 1991; Maldonado et al., 2006; Ghosh et al., 2014; Liu et al., 2017; Moghe et al., 2017). These compounds serve as microbial and herbivory defense agents (Goffreda et al., 1989; Leckie et al., 2016; Luu et al., 2017; Smeda et al., 2018) and contain a sugar core esterified to multiple acyl chains (Fan et al., 2019). The acyl chains are branched or straight and are proposed to be derived from amino acid or fatty acid metabolism, respectively (Walters and Steffens, 1990; Kroumova and Wagner, 2009; Ning et al., 2015). Acylsugar cores from Solanaceae species are primarily sucrose or sometimes glucose (King et al., 1986; King and Calhoun, 1988; Maldonado et al., 2006; Ghosh et al., 2014; Liu et al., 2017; Moghe et al., 2017; Leong et al., 2019). *Solanum lanceolatum* is a published exception, accumulating acylated disaccharide cores that contain *myo*-inositol bound to glucose or xylose (Herrera-Salgado et al., 2005). Moghe and colleagues screened acylsugars in plants across the Solanaceae family (Moghe et al., 2017). One species – the South American fruit crop *Solanum quitoense* – produced unusual acylsugars that yielded lower signals of fragment ions generated in a mass spectrometer than acylsucroses or acylglucoses from other members of the Solanaceae (Hurney, 2018).

Acylsugar acyltransferases (ASATs) are clade III BAHD (BEAT, AHCT, HCT, and DAT) acyltransferase family enzymes (D’Auria, 2006; Moghe et al., 2017), which catalyze the core acylation reactions of characterized acylsugar biosynthetic pathways (Schilmiller et al., 2012a; Schilmiller et al., 2015; Fan et al., 2016; Moghe et al., 2017; Nadakuduti et al., 2017; Fan et al., 2019). ASATs sequentially transfer acyl chains from acyl-CoAs to generate acylsucroses in plants as phylogenetically divergent as *Solanum lycopersicum, Solanum pennellii, Petunia axillaris*, and *Salpiglossis sinuata* (Schilmiller et al., 2012a; Schilmiller et al., 2015; Fan et al., 2016; Fan et al., 2017; Moghe et al., 2017; Nadakuduti et al., 2017). Expression of characterized ASATs is enriched in glandular trichomes relative to non-acylsugar producing tissues (Ning et al., 2015; Moghe et al., 2017; Nadakuduti et al., 2017; Leong et al., 2019; Mandal et al., 2020). Phylogenetic and functional analyses revealed that orthologs have similar – but distinct – substrate selectivity in acylsugar biosynthesis across the Solanaceae family (Moghe et al., 2017; Nadakuduti et al., 2017). The combination of tissue enrichment and phylogenetic relatedness are powerful tools to identify enzymes in other acylsugar pathways.

Here we report structural characterization of monosaccharide acylsugars that are built on a *myo*-inositol core from *S. quitoense* aerial tissues, along with demonstration of function of a triacylinositol acetyltransferase (TAIAT) that catalyzes the fourth and final acylation step in *S. quitoense* acylsugar biosynthesis. Liquid chromatography–mass spectrometry (LC-MS) and nuclear magnetic resonance (NMR) spectroscopy analysis revealed that these are *myo*-acylinositols with three to four acyl chains of two, ten and twelve carbon atoms as major acylsugars. We used RNAseq data and phylogenetic analysis to identify a protein related to Sl-ASAT4 – the enzyme that acetylates triacylated sucroses in *S. lycopersicum*. A combination of virus-induced gene silencing (VIGS) and *in vitro* biochemistry demonstrated this enzyme acetylates *S. quitoense* triacylinositols at the 4-position of *myo*-inositol. This analysis reveals parallels between the *S. quitoense* acylinositol and *S. lycopersicum* acylsucrose biosynthetic pathways that suggest a common recent evolutionary origin.

## Results

### NMR and LC-MS analysis reveal *Solanum quitoense* acylinositol structures

LC-MS analysis of *Solanum quitoense* leaf surface extracts revealed previously-undescribed acylhexoses and acyldisaccharides (**Fig. 1A**). Subsequent generation of MS-MS product ions of [M+formate]^-^ ions of the four major acylhexoses yielded spectra dominated by fragment ions assigned as fatty acid carboxylates of ten (*m/z* 171.14) and twelve (*m/z* 199.17) carbons, but few other fragments (**Supplemental fig. S1**). Lack of intermediate sugar core-containing fragment ions derived from neutral losses of acyl groups is inconsistent with previous collision-induced dissociation (CID) mass spectra observed for acylsucroses and acylglucoses analyzed (Leong et al., 2019). In contrast, MS-MS product ion mass spectra of [M+NH_4_]^+^ of the four *S. quitoense* acylhexoses yielded fragment ions consistent with neutral losses of two, ten or twelve carbon aliphatic acids and ketenes (**Supplemental fig. S2**). The fatty acid ions and neutral losses from these compounds indicated that these acylsugars have 1-2 two-carbon acyl chains and two longer chains of ten or twelve carbons on hexose and disaccharide cores.

**Figure 1.**
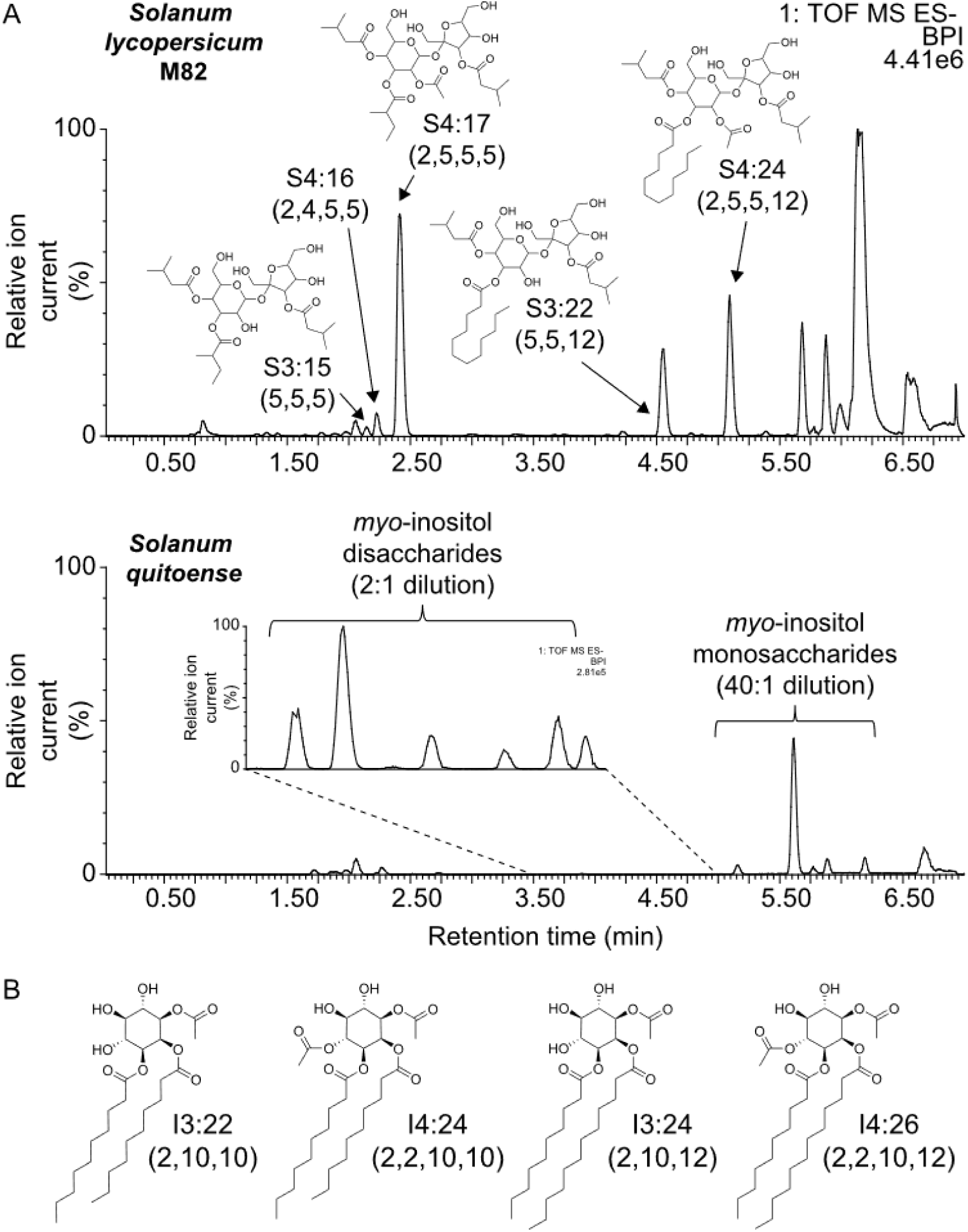
Profiling and structures of *Solanum quitoense* acylsugars. **(A)** Comparison of base peak intensity LC-MS chromatograms collected using ESI^-^ mode of *Solanum lycopersicum* and *Solanum quitoense* acylsugars. **(B)** Nuclear magnetic resonance-derived structures of tri- and tetraacylated *myo*-inositol monosaccharides purified from *S. quitoense*. Note: NMR analysis was unable to determine the exact location of nC10 and nC12 acyl chains in I3:24 (2,10,12) and I4:26 (2,2,10,12). Therefore, the positions containing medium length acyl chains are correct, but nC10 and nC12 chains may be reversed.

NMR analysis of the purified compounds – the four highest abundance *S. quitoense* acylsugars – confirmed the hypothesis that the sugar core differed from glucose. These sugar esters each contain three to four acyl chains – two unbranched and saturated C10 or C12 acyl chains, and one or two acetyl (C2) groups consistent with the LC-MS-MS analysis (**Fig. 1B, Supplemental fig. S2, Supplemental Table S1**). Each metabolite contains a *myo*-inositol core (**Fig. 1B, Supplemental Table S1**), which is a sugar alcohol acylsugar core previously reported in one species in the Solanaceae family (Herrera-Salgado et al., 2005). There are five less abundant acylinositols modified by glycosylation with glucose, xylose, or N-acetylglucosamine (Hurney, 2018). Our study focused on the non-glycosylated acylinositols, representing >85% of the integrated LC-MS acylsugar peak areas (Hurney, 2018). The nomenclature used to describe these compounds is as follows: IX:Y (A,B,C,D), where ‘I’ signifies the *myo*-inositol core, ‘X’ represents the total number of acyl esters, ‘Y’ is the sum of carbons in those acyl chains, while ‘A’, ‘B’, ‘C’ and ‘D’ represent the number of carbons in each acyl chain. Taken together, the NMR results revealed the four most abundant *S. quitoense* acylsugars as: I3:22 (2,10,10); I4:24 (2,2,10,10); I3:24 (2,10,12); I4:26 (2,2,10,12) (**Fig. 1, Supplemental Table S1**).

### Identification of acylinositol acyltransferase candidates in *S. quitoense*

With knowledge of acylinositol structures in hand, we sought candidates for their biosynthesis, using several criteria to analyze RNAseq data from Moghe et al. (2017). First, we selected candidate BAHD enzymes based on the presence of two conserved BAHD signatures motifs (D’Auria, 2006): a HXXXD catalytic motif and a DFGWG structural motif, both present in a specific orientation. A second criterion was that candidates had to be between 400-500 amino acids in length, as found in characterized functional BAHDs. Together, these steps resulted in a list of 42 candidates. Previously characterized ASATs are expressed and enriched in glandular trichomes relative to non-acylsugar accumulating tissues (Schilmiller et al., 2012a; Ning et al., 2015; Fan et al., 2016; Moghe et al., 2017; Nadakuduti et al., 2017; Leong et al., 2019; Mandal et al., 2020). Thus, the 17 candidates with >500 reads in the trichome samples were selected for further analysis.

We hypothesized that the acylinositol pathway is evolutionarily related to acylsucrose biosynthesis (Fan et al., 2019), selecting the six putative BAHDs most closely related to previously characterized ASATs in clade III of BAHD acyltransferases (**Fig. 2, Supplemental fig. S3**)(Schilmiller et al., 2012a; Schilmiller et al., 2015; Fan et al., 2016; Moghe et al., 2017; Nadakuduti et al., 2017). Given the presence of C2 groups on the NMR-characterized acylinositols, the following candidates were of great interest; three transcripts – c38687_g1_i4, c38687_g1_i1, and c38687_g2_i1 (c#####_g#_i# represents distinct transcripts in the RNAseq assembly) – were identified as the closest homologs of Sl-ASAT4, which catalyzes acetylation of triacylsucroses in cultivated tomato (**Table 1, Fig. 2, Fig. 3A, Supplemental fig. S3**). Our analysis focused on c38687_g1_i4 because it has the 7^th^ largest number of reads of all transcripts in the trichome dataset and is enriched in trichomes relative to stem tissue (**Table 1**). Results of the *in planta* and biochemical analyses described below lead to the conclusion that this gene is a triacylinositol acetyltransferase (TAIAT).

**Table 1.** RNAseq data for *S. quitoense* ASAT4 homologs. Data are derived from Moghe et al. (Moghe et al., 2017).

| Transcript name | Average trichome reads | Average petiole reads |
| --- | --- | --- |
| c38687_g1_i1 | 1634 | 3 |
| c38687_g1_i4 (TAIAT) | 47432 | 143 |
| c38687_g2_i1 | 2679 | 11 |

**Figure 2.**
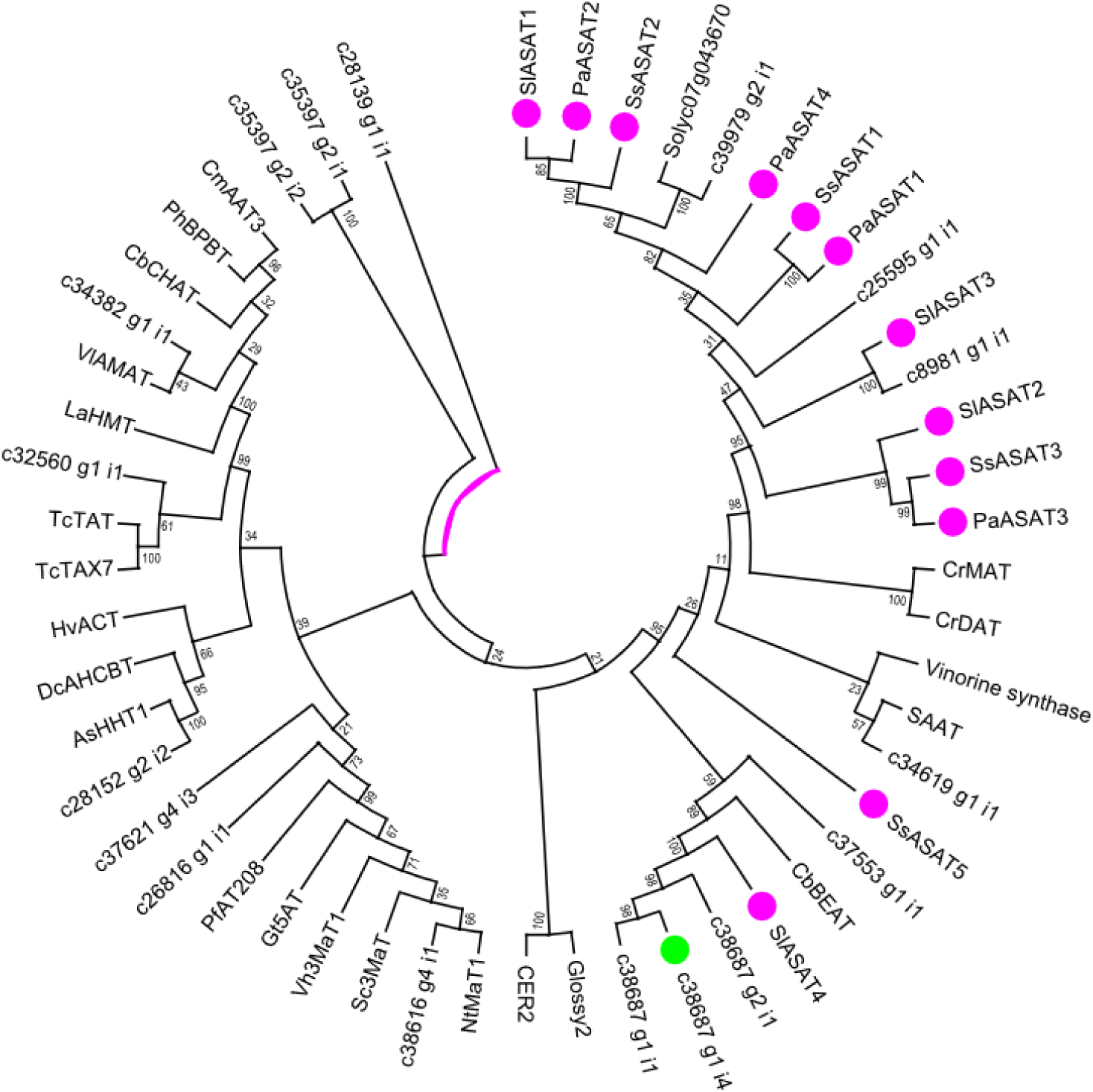
Phylogenetic analysis of ASAT candidates and previously characterized BAHDs. Phylogenetic tree showing several previously characterized ASATs and other BAHD acyltransferases. Previously characterized ASATs are marked by magenta circles, while TAIAT is marked by a green circle. Protein sequences were aligned using MUSCLE with default parameters in MEGA X. Maximum likelihood method was used to perform the phylogenetic analysis. Jones-Taylor-Thornton (JTT) +G with five rate categories with 1000 bootstrap replicates using partial deletion (30% gaps) for tree reconstruction was used. Further details are included in Materials and Methods.

**Figure 3.**
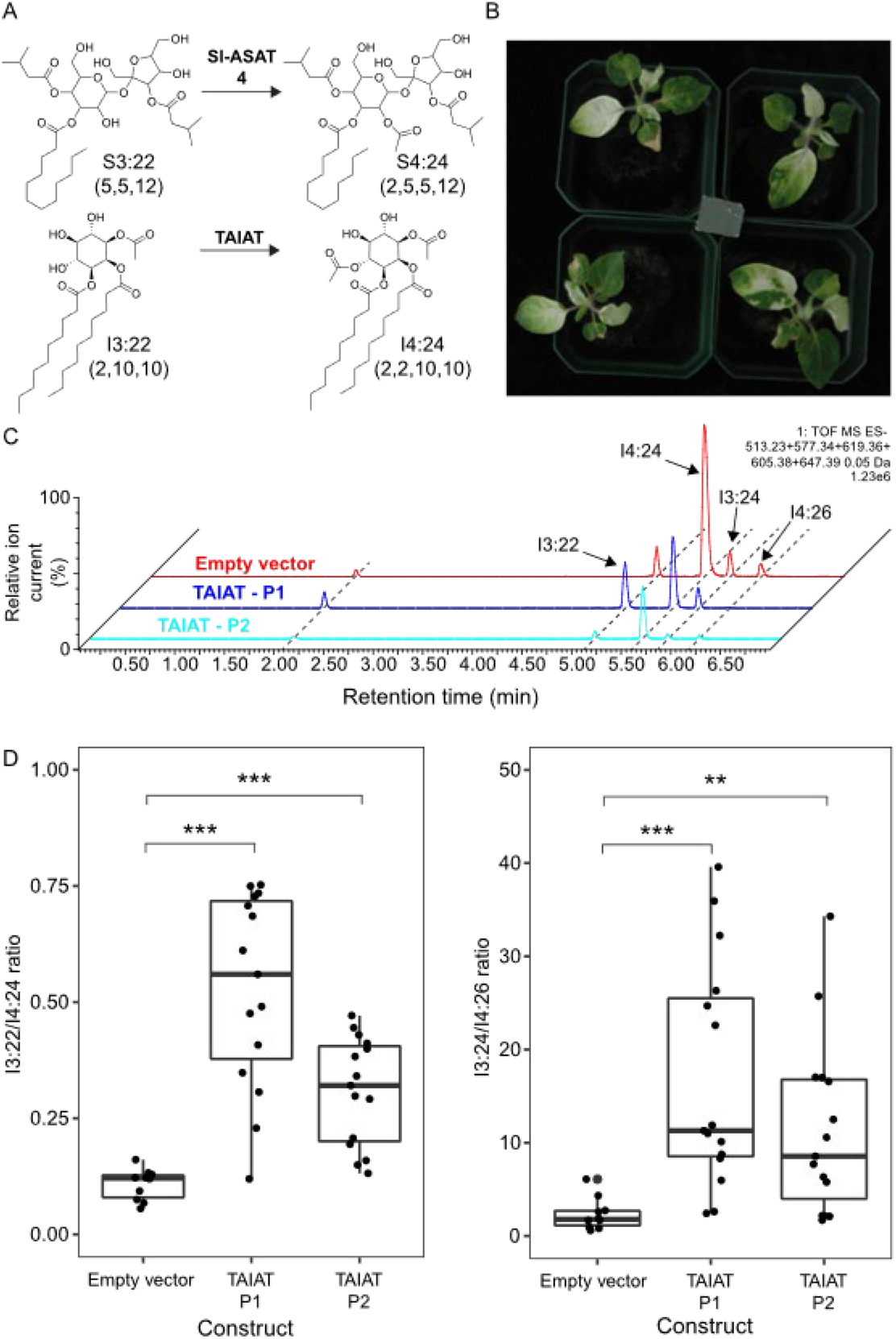
Virus-induced gene silencing (VIGS) of TAIAT in *S. quitoense*. **(A)** Characterized or proposed enzymatic activity of Sl-ASAT4 or TAIAT, respectively. **(B)** Phenotype of phytoene desaturase (PDS)-silenced plants that were used as a visual marker to evaluate VIGS efficacy. Pot length and width is uniform at 2.875 inches. **(C)** Acylsugar analysis of TAIAT-targeted and empty vector VIGS plants. Acylsugars were analyzed using LC-MS in ESI- mode. A combined extracted ion chromatogram is displayed showing: telmisartan (internal standard), (*m/z*: 513.23); I3:22 (2,10,10), (*m/z*: 577.34); I4:24 (2,2,10,10), (*m/z*: 619.36); I3:24 (2,10,12), (*m/z*: 605.38); I4:26 (2,2,10,12), (*m/z*: 647.39). LC-MS extracts were from similar size leaves (≤ 0.15 mg difference between leaves). TAIAT-targeted lines are blue or cyan, while the empty vector plants are red. **(D)** Comparison of total tri- and tetraacylinositol LC-MS peak areas in TAIAT-targeted and empty vector VIGS plants. Acylsugars were analyzed as in C. Whiskers represent minimum and maximum values less than 1.5 times the interquartile range from the first and third quartiles, respectively. Expression values outside this range are represented by dots. P-values: ** p < 0.01; *** p < 0.001. Welch’s two sample t-test was used. TAIAT-targeted plants (n=15), Empty vector plants (n=10).

### *In vivo* analysis of TAIAT function

The acylinositol NMR structures are consistent with the hypothesis that monoacetylated *S. quitoense* triacylinositols with a single acetyl ester are biosynthetic precursors of tetraacylinositols with two acetyl esters (**Fig. 1, Supplemental Table S1**). This predicts that acetylation of the 4-position hydroxyl is the final step in tetraacylated inositol biosynthesis because both I3:22 (2,10,10)/I4:24 (2,2,10,10) and I3:24 (2,10,12)/I4:26 (2,2,10,12) differ by acetylation at this position (**Fig. 1, Fig. 3A**).

Virus-induced gene silencing (VIGS) was developed for *S. quitoense* (**Fig. 3B**) to test whether TAIAT has a role in acylinositol biosynthesis. As predicted for an enzyme catalyzing the second acetylation reaction, VIGS plants derived from two different TAIAT constructs showed strong and statistically significant increases in the ratio of monoacetylated I3:22 (2,10,10) to diacetylated I4:24 (2,2,10,10) relative to the controls (**Fig. 3C and D, Supplemental Table S2**). A similar increase in the ratio of I3:24 (2,10,12) was observed relative to I4:26 (2,2,10,12) (**Fig. 3C, D, Supplemental Table S2**). qPCR analysis revealed reductions in TAIAT expression in some of the target plants relative to the controls (**Supplemental fig. S4**). The increased ratio of triacylated- to tetraacylated-inositols in TAIAT VIGS plants supports the hypothesis that I3:22 (2,10,10) and I3:24 (2,10,12) are direct precursors of the more acetylated products and that TAIAT catalyzes the acetylation of these triacylinositols.

### *In vitro* analysis of TAIAT

We tested TAIAT *in vitro* activity to determine whether it can acetylate I3:22 (2,10,10) and I3:24 (2,10,12). His-tagged TAIAT was expressed in *Escherichia coli* and purified using Ni-Nitrilotriacetic acid (Ni-NTA). Acylinositol substrates were enriched from *S. quitoense* leaf surface extracts using semi-preparative liquid chromatography.

*In vitro* assays validated that TAIAT acetylates *S. quitoense* triacylinositols. TAIAT acetylated I3:22 (2,10,10) to form I4:24 (2,2,10,10) (**Fig. 4A**) and also converted I3:24 (2,10,12) to I4:26 (2,2,10,12) (**Fig. 4B**). *In vitro* products and *in planta* acylsugars co-eluted with – and fragmented similarly to – each other (**Supplemental fig. S5, Supplemental fig. S6**). The two other ASAT4 homologs show minimal activity with triacylinositols (< 1% of TAIAT product accumulation) (**Supplemental fig. S7, Supplemental fig. S8**). These results indicate that TAIAT converts tri- to tetraacylinositols in *S. quitoense*.

**Figure 4.**
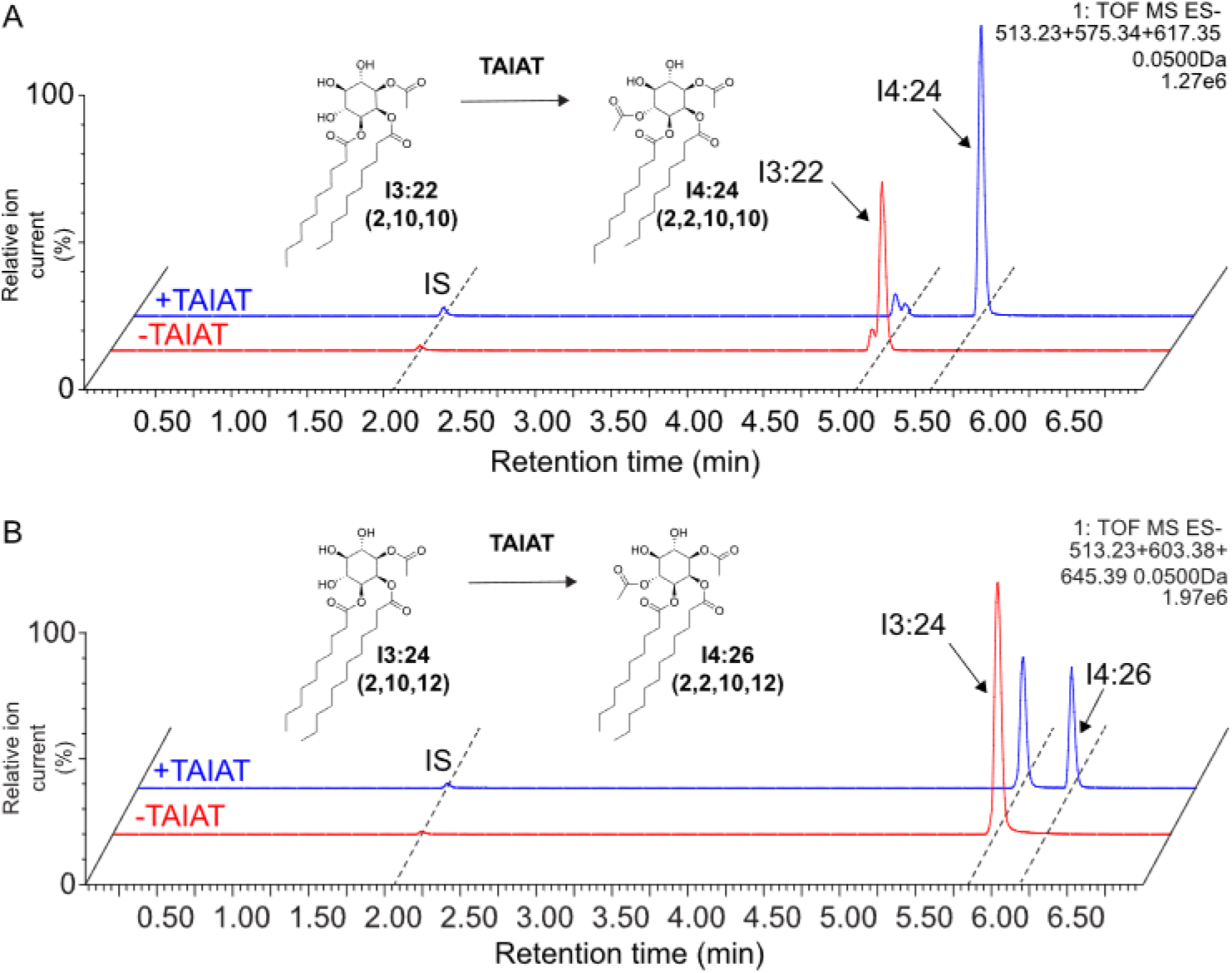
*In vitro* assays using TAIAT on plant-derived triacylinositols. **(A)** LC-MS analysis of *in vitro* assay products provide evidence that TAIAT acetylates I3:22 (2,10,10). Combined extracted ion chromatogram (ESI^-^) showing telmisartan internal standard, (*m/z*: 513.23); I3:22 (2,10,10), (*m/z*: 575.34); I4:24 (2,2,10,10), (*m/z*: 617.35). **(B)** LC-MS analysis of *in vitro* assay products provide evidence that TAIAT acetylates I3:24 (2,10,12). Combined extracted ion chromatogram showing telmisartan internal standard, (*m/z*: 513.23); I3:24 (2,10,12), (*m/z*: 603.38); and I4:26 (2,2,10,12), (*m/z*: 645.39). Acylsugars were detected as formate adducts using ESI^-^ mode. The blue traces represent the full TAIAT assay, whereas the red traces have water substituted for TAIAT.

We took advantage of the ability of BAHDs to catalyze reverse reactions when provided with its usual product and coenzyme A, to ask whether TAIAT specifically catalyzes transfer of the acetyl group from position 4. Reactions were performed using extracts containing individual *in planta* acylinositols that had two acetyl groups. TAIAT deacetylated both I4:24 (2,2,10,10) and I4:26 (2,2,10,12) to generate single chromatographic peaks for I3:22 (2,10,10) and I3:24 (2,10,12) respectively, using two different LC columns (**Supplemental fig. S9**). The reaction products co-eluted with – and fragmented similarly to – *S. quitoense* triacylinositols (**Supplemental fig. S6, Supplemental fig. S10**). The combination of forward and reverse *in vitro* enzymatic activities and *in planta* VIGS phenotypes provide compelling evidence that TAIAT acetylates *S. quitoense* triacylinositols.

## Discussion

Acylsugars constitute a diverse group of specialized metabolites produced across the Solanaceae family (Moghe et al., 2017). Variation in acyl chain length, branching pattern or occasionally other constituents such as malonyl groups all contribute to this diversity (Fan et al., 2019). Other factors include differences in acylation position, acyl-CoA specificity, and sugar core diversity (Fan et al., 2016; Fan et al., 2017; Moghe et al., 2017; Fan et al., 2019). Most characterized acylsugar-producing species accumulate primarily acylsucroses with isolated occurrences of acylglucoses in *Nicotiana miersii, Datura metel* or *S. pennellii* (Burke et al., 1987; King and Calhoun, 1988; Matsuzaki et al., 1989; Ghosh et al., 2014; Liu et al., 2017; Moghe et al., 2017). Accumulation of acylsugars has only been explored in a small fraction of the approximately 2700 known species in the Solanaceae. Our study focused on one such underexplored species, *S. quitoense*.

The structures of *S. quitoense* acylsugars we characterized are distinct from those previously reported in two ways. First, these metabolites represent one of the first examples of *myo*-inositol in acylsugars: the other published case is *Solanum lanceolatum* (Herrera-Salgado et al., 2005) where inositol was a component of disaccharide cores. Acyl chain makeup is another differentiating element with exclusively C10-C12 unbranched acyl chains and acetyl groups.

This contrasts with *S. lycopersicum* and *S. sinuata*, which produce acylsugars with branched acyl chains presumably derived from branched chain amino acid precursors (Walters and Steffens, 1990; Kroumova and Wagner, 2009; Ghosh et al., 2014; Ning et al., 2015; Liu et al., 2017; Moghe et al., 2017). The unusual sugar core and acyl chain structures led us to characterize acylinositol biosynthesis.

We characterized the BAHD acetyltransferase, TAIAT, as the enzyme that catalyzes a terminal step in *S. quitoense* tetraacylinositol biosynthesis. Two lines of evidence indicate that TAIAT acetylates *S. quitoense* triacylinositols. Use of a newly established *S. quitoense* VIGS protocol revealed that reduction in TAIAT expression caused increased ratios of tri- to tetraacylated inositols (**Fig. 3**). *In vitro* assays demonstrated that TAIAT can acetylate *S. quitoense* produced I3:22 (2,10,10) and I3:24 (2,2,10,12), and – in the reverse reaction – deacetylate I4:24 (2,2,10,10) and I4:26 (2,2,12,12) (**Fig. 4, Supplemental fig. S9**). The combined *in planta* and *in vitro* findings implicate TAIAT as the last acetylation step in tetraacylinositol biosynthesis.

TAIAT substrates are similar to Sl-ASAT4. Our results show that TAIAT uses acetyl-CoA – the acyl donor substrate of several BAHD enzymes in Clade III including Sl-ASAT4 (**Fig. 4**) (D’Auria, 2006; Schilmiller et al., 2012a; Moghe et al., 2017). The acyl acceptor substrates are also similar: *S. lycopersicum* Sl-ASAT4 acetylates triacylsucroses whereas TAIAT acetylates triacylinositols in *S. quitoense*.

The combination of trichome-enriched expression and phylogenetic analysis is a powerful tool to identify uncharacterized BAHDs involved in acylsugar biosynthesis (Moghe et al., 2017; Nadakuduti et al., 2017). *S. quitoense* trichome RNAseq data contain five putative ASAT homologs in addition to TAIAT (**Fig. 2**). It is possible that these homologs play a role in one or more of the remaining biosynthetic steps based on their trichome expression and phylogenetic relationships.

Our study uncovered parallels that suggest a common recent evolutionary origin between acylinositol and acylsucrose biosynthesis. The acetyl-CoA substrate used by Sl-ASAT4 and TAIAT, their close phylogenetic relationship, and enrichment in trichomes are evidence for a link between these pathways. Phylogenetic analysis of acylsugar-producing species point to sucrose as the ASAT ancestral acyl acceptor – it is the sugar acceptor in earlier diverging lineages leading to *S. sinuata* and *P. axillaris* (Moghe et al., 2017; Nadakuduti et al., 2017). Further, acylsucrose-accumulating species are present throughout the Solanaceae family (Burke et al., 1987; King and Calhoun, 1988; Matsuzaki et al., 1991; Maldonado et al., 2006; Ghosh et al., 2014; Liu et al., 2017; Moghe et al., 2017), whereas reported acylinositol accumulators are *S. quitoense* and *S. lanceolatum* (**Fig. 1**) (Herrera-Salgado et al., 2005). Both species are members of the Leptostemonum or ‘Spiny Solanum’ clade. These limited data suggest that acylinositol biosynthesis more recently emerged from the acylsucrose pathway; screening the Leptostenomum clade and the *Solanum* genus more broadly would provide insight into when acylinositol biosynthesis emerged. Acylinositol biosynthesis provides opportunities to better understand acylsugar diversification and broaden our knowledge of both pathway and enzyme evolution.

## Materials and Methods

### Heterologous protein expression and purification from *Escherichia coli*

Heterologous protein expression was achieved using pET28b(+) (EMD Millipore, Burlington, MA), in which open reading frames for the enzymes were cloned into BamHI/XhoI doubly digested vector. Inserts were amplified by PCR using Q5 2x Hotstart mastermix (NEB, Ipswich, MA), and purified by agarose gel purification and extraction (primers: TAIAT, c38687_g1_F/R; c38687_g1_i1, c38687_g1_F/c38687_g1_i1_R; c38687_g2_i1, c38687_g2_F/R). The doubly-digested vectors were assembled with a single fragment containing the ORF containing 5’ and 3’ adapters for Gibson assembly using 2x NEB Hifi Mastermix (NEB, Ipswich, MA). The finished constructs were transformed into BL21 Rosetta (DE3) cells (EMD Millipore, Burlington, MA) and verified using colony PCR and Sanger sequenced using T7 promoter and terminator primers. Primer sequences are listed in Supplemental Table S3.

LB overnight cultures with kanamycin (50 µg/mL) and chloramphenicol (33 µg/mL) were inoculated with a single colony of the bacterial strain containing the desired construct and incubated at 37 °C, 225 rpm, overnight. Larger LB cultures (1L) were inoculated 500:1 with the same antibiotics and incubated at the same temperature and speed. OD600 of the cultures was monitored until between 0.5 and 0.8. Cultures were chilled on ice for 15 minutes, at which point, IPTG was added to a final concentration of 50 µM. Cultures were incubated at 16 °C, 180 rpm for 16 hours.

Note: All of the following steps were on ice or at 4°C.

Cultures were centrifuged at 4,000g for 10 minutes to collect the cells and repeated until all the culture was processed (4 °C). The cell pellets were resuspended in 25 mL of extraction buffer (50 mM NaPO_4_, 300 mM NaCl, 20 mM imidazole, 5 mM 2-mercaptoethanol, pH 8.0) by vortexing. The cell suspension was sonicated for 8 cycles (30 seconds on, intensity 4, 30 seconds on ice). The cellular extracts were centrifuged at 30,000g for 10 minutes. The supernatant was transferred into another tube and centrifuged again at the same speed and duration. Ni-NTA resin (Qiagen, Hilden, Germany) was centrifuged at 1,000g for 1 minute, resuspended in 1 mL of extraction buffer. The slurry was centrifuged again at 1,000g for 1 minute and the supernatant was decanted. The resin was resuspended using the centrifuged extract and incubated at 4 °C, nutating for 1 hour. The slurry was centrifuged at 3,200g for 5 minutes, and supernatant decanted. The resin was resuspended in 5 mL of extraction buffer and transferred to a gravity flow column (Biorad, Hercules, CA). After loading, the resin was washed with 3 column volumes of extraction buffer (∼30 mL). The resin was further washed with 1 column volume of wash buffer (extraction buffer with 40 mM imidazole). The remaining protein was eluted and collected using 2 mL of elution buffer after a 1 minute incubation with the resin. The elutate was diluted into 15 mL of storage buffer (extraction buffer, but no imidazole) and concentrated using 10 kDa centrifugal filter units (EMD Millipore, Burlington, MA), and repeated until diluted 1,000 fold. An equal volume of 80% glycerol was added to the elution, mixed, and stored at -20 °C. SDS-PAGE was used to analyzed elution fractions, and the presence of enzymes were confirmed by immunoblot using the anti-His6 antibody conjugated to peroxidase (BMG–His-1 monoclonal antibody) (Roche, Mannheim, Germany).

### Enzyme assays

Assays were run in 100 mM sodium phosphate buffer at a total volume of 60 µL with pH 6.0 as the default unless otherwise stated. Acyl-CoA (or coenzyme A) was added to a final concentration of 100 µM and obtained from Sigma-Aldrich (Sigma-Aldrich, St. Louis, MO). Acylsugar acceptors – such as I3:22 – were dried down using a speed vac and dissolved in an ethanol:water mixture (1:1) with 1 µL added to the reaction. 6 µL of enzyme was added to each reaction (or water). The assays were incubated at 30 °C for 30 minutes unless otherwise stated. After the incubation, 2 volumes of stop solution – composed of a 1:1 of acetonitrile and isopropanol with 0.1% formic acid and 1 µM telmisartan as internal standard (Sigma-Aldrich, St. Louis, MO) – were added to the assays and mixed by pipetting. Reactions were stored in the -20 °C freezer for 20 minutes and centrifuged at 17,000g for 5 minutes. The supernatant was transferred to LC-MS autosampler vials and stored at -20 °C. For negative control assays in **Supplemental fig. S7 and 8**, an aliquot of the enzyme was incubated at 95°C for 10 minutes to inactivate it.

### LC-MS analysis

LC-MS analyses of products from enzyme assays and extracts of plant tissues were analyzed on a Waters Acquity UPLC coupled to a Waters Xevo G2-XS QToF mass spectrometer (Waters Corporation, Milford, MA). 10 µL of the extracts were injected into an Ascentis Express C18 HPLC column (10 cm x 2.1 mm, 2.7 µm) or Ascentis Express F5 HPLC column (10 cm x 2.1 mm, 2.7 µm) (Sigma-Aldrich, St. Louis, MO) that was maintained at 40 °C. The LC-MS methods used the following solvents: 10 mM aqueous ammonium formate, adjusted to pH 2.8 with formic acid as solvent A, and 100% acetonitrile as solvent B. A flow rate of 0.3 mL/minute was used unless otherwise specified. A 7-minute linear elution gradient consisted of 5% B at 0 minutes, 60% B at 1 minute, 100% B at 5 minutes, held at 100% B until 6 minutes, 5% B at 6.01 minutes and held at 5% until 7 minutes. A 21-minute linear elution gradient consisted of 5% B at 0 minutes, 60% B at 3 minutes, 100% B at 15 minutes, held at 100% B until 18 minutes, 5% B at 18.01 minutes and held at 5% until 21 minutes.

For ESI^-^ MS settings: capillary voltage, 2.00 kV; source temperature, 100 °C; desolvation temperature, 350 °C; desolvation nitrogen gas flow rate, 600 liters/hour; cone voltage, 40 V; mass range, *m/z* 50-1000 (with spectra accumulated at 0.1s per function). Three acquisition functions were used to acquire spectra at different collision energies (0, 15, and 35 V). Lock mass correction was performed using leucine enkephalin [M-H]^-^ as the reference and corrections were applied during data acquisition.

For ESI^+^ MS settings: capillary voltage, 3.00 kV; source temperature, 100 °C; desolvation temperature, 350 °C; desolvation nitrogen gas flow rate, 600 liters/hour; cone voltage, 35 V; mass range, *m/z* 50-1000 (with spectra accumulated at 0.1 s per function). Two acquisition functions were used to acquire spectra at different collision energy settings (0, 10-60 V ramp). Lock mass correction was performed using leucine enkephalin [M+H]^+^ as the reference and corrections were applied during data acquisition.

Data-dependent LC-MS-MS analyses were performed on a LC-20ADvp ternary pump (Shimadzu) coupled via an Ascentis Express C18 column (2.1 x 100 mm, 2.7 µm, Supelco) to a Xevo G2-XS mass spectrometer (Waters). Ion source parameters were as described above for LC/MS analyses. Column temperature was held at 50°C. Gradient elution employed solvent A (10 mM aqueous ammonium formate, adjusted to pH 3.0 with formic acid) and solvent B (acetonitrile). A linear gradient (A/B) was as follows: 0-1 min (99/1), 32 min (0/100), 36 min (0/100), 36.01 min (99/1), 40 min (99/1). Survey scans were acquired over *m/z* 50-1000 with 0.2 seconds/scan. The three most abundant ions in the survey scan were selected for MS/MS product ion spectra at 0.25 s/scan, using a collision voltage of 20 V in negative-ion and positive-ion modes.

### Acylsugar purification for enzyme assays

Acylsugars were extracted from 20 leaves from 12-week old *S. quitoense* plants into 500 mL of methanol w/ 0.1% formic acid with gentle agitation in a 1L beaker. The methanol was transferred to a round-bottom flask and evaporated under reduced pressure using a rotary evaporator with a warm water bath (∼40 °C). The dried residue was dissolved in 2 mL of acetonitrile (0.1% formic acid) and stored in an LC-MS vial at -20°C. 500 µL of the solution was transferred into a microcentrifuge tube and evaporated to dryness using a speed vac. The residue was dissolved in 550 µL of 4:1 acetonitrile:water w/ 0.1% formic acid. Five 100-µL injections were made onto a Waters 2795 HPLC equipped with a C18 semipreparative column (Acclaim C18 5 µm, 120 Å, 4.6 x 150 mm) using a 63-minute chromatographic method to separate the acylsugars with a flow rate of 1.5 mL/minute and column temperature of 30 °C. Solvent A was water w/ 0.1% formic acid, and Solvent B was acetonitrile.

The chromatographic gradient is as follows: 5% B at 0.00 minutes, 60% B at 1.00 minute, 100% B at 50.00 minutes, hold at 100% B until 60.00 minutes, 5% B at 60.01 minutes, and hold at 5% B until 63 minutes. 1-minute fractions were collected for a total of 63 fractions. Fractions were screened (LC-MS) for the presence of acylsugars, and fractions judged to have sufficient purity were pooled and evaporated to dryness in the speed vac. Each acylsugar was resuspended in acetonitrile with 0.1% formic acid, transferred to LC-MS vials, and stored in the - 20 °C freezer. 200-µL aliquots of the acylsugars were transferred to microcentrifuge tubes with glass inserts and evaporated to dryness using a speed vac before dissolution in 20 µL of 1:1 ethanol:water mixture for use in enzyme assays.

### Gene identification and phylogenetic analysis

All transcript assemblies are from Moghe et al. (2017) and were analyzed using Geneious R8.1.9. Sequences were selected by HXXXD motifs, which were detected using the ‘Search for Motif’ function (0 mismatches). Those sequences were further parsed to include only those that contain the DFGWG motif (1 mismatch) using the same function. The remaining sequences were further screened to a length of 400-500 amino acids, and by the relative positions of the two motifs (D’Auria, 2006). Sequences were aligned against several other BAHD sequences from D’Auria (2006), and several characterized ASATs (Schilmiller et al., 2012a; Schilmiller et al., 2015; Fan et al., 2016; Moghe et al., 2017). Both transcript and expression data are available in (Moghe et al., 2017).

Phylogenetic reconstructions were performed using MEGA X (Kumar et al., 2018). For phylogenetic reconstructions, amino acid sequences were aligned using MUSCLE under default parameters. A maximum likelihood method was used to generate the phylogenetic tree. The model selection feature in MEGA X was used to determine the best evolutionary model for the maximum likelihood method – Jones-Taylor-Thornton (JTT)+G with five rate categories. 1000 bootstrap replicates were performed using partial deletion (30% gaps) for tree reconstruction.

For multiple sequence alignments and amino acid identities present in figures, alignments were performed in Geneious R8.1.9 using the MUSCLE algorithm under default settings.

### Plant growth conditions

Plants were grown at 24°C, 16-hour light/8-hour dark cycle with a light intensity of 70 µmol m^-2^ s^-2^ PPFD in Jiffy 7 peat pellets (Jiffy, Kristiansand, Norway). Plants were watered four times per week with deionized water and fertilized once per week with half-strength Hoagland’s solution. These were the plant growth conditions unless otherwise specified.

### Acylsugar analysis

The interactive protocol for acylsugar extracts is available at Protocols.io at: https://dx.doi.org/10.17504/protocols.io.xj2fkqe The acylsugar extraction protocol was described in (Leong et al., 2019). LC-MS conditions used for acylsugar analysis were described in the LC-MS analysis section. Acylsugar sugars were analyzed in ESI^-^ as formate adducts and in ESI^+^ mode as ammonium adducts.

### VIGS analysis

pTRV2-LIC was digested using PstI-HF to generate the linearized vector. Fragments were amplified using PCR with adapters for ligation into pTRV2-LIC (c38687_g1_p1_F/R, c38687_g1_p2_F/R, or SqPDS_VIGS_F/R). Note: c38687_g1_p1_R is located in the 3’ UTR. Both the linearized vector and PCR products were purified using a 1% agarose gel and gel extracted using an Omega EZNA gel extraction kit. Both the PCR fragment and the linearized vector were incubated in separate 5 µL reactions using NEB 2.1 as buffer with T4 DNA polymerase and 5 mM dATP or dTTP (PCR insert/Vector). The reactions were incubated at 22 °C for 30 minutes, subsequently incubated at 70 °C for 20 minutes. The reactions were then stored on ice. 1 µL of the pTRV2-LIC reaction and 2 µL of the PCR-LIC reaction were mixed by pipetting. Reactions were incubated at 65 °C for 2 minutes, then 22 °C for 10 minutes. After which the constructs were transformed into chemically competent *E.coli* DH5α cells. Primer sequences available in Supplemental Table S3.

Constructs were tested for the presence of the insert using colony PCR and pTRV2_LIC_seq_F/R primers showing a 300 bp insertion. Positive constructs were miniprepped (Qiagen, Hilden, Germany) and sanger sequenced using the same primers. Sequenced constructs and pTRV1 were transformed into agrobacterium strain, GV3101, using the protocol described previously except on LB plates with kanamycin (50 µg/mL), rifampicin (50 µg/mL), and gentamycin (10 µg/mL) (Leong et al., 2019). Colonies were assayed for the presence of the insert using the colony PCR and the pTRV2_LIC_seq_F/R primers previously described. The presence of the pTRV1 vector in GV3101 was assayed using colony PCR primers, pTRV1_F/R. Primer sequences available in Supplemental Table S3.

The VIGS protocol was adapted from (Velásquez et al., 2009). Seeds were incubated in 10% bleach for 30 minutes, followed by 5-6 washes with water. Seeds were transferred to a Petri dish with Whatman paper and water in the bottom of the dish. Seeds were stored in the dark at room temperature until hypocotyl emergence, at which point they were moved to a window sill. Once cotyledons emerged, seedlings were transferred to peat pots (Jiffy, Kristiansand, Norway), and grown for approximately 1 week under 16/8 day/night cycle at 24 °C until inoculated. At 2 days pre-inoculation, LB cultures (Kan/Rif/Gent) were inoculated with the cultures used for leaf inoculation. The strains have constructs containing the gene of interest (GOI) in pTRV2-LIC, an empty vector pTRV2-LIC, and pTRV1. Cultures were grown overnight at 30 °C with shaking at 225 rpm. Larger cultures composed of induction media (4.88g MES, 2.5g glucose, 0.12g sodium phosphate monobasic monohydrate in 500 mL, pH 5.6, 200 µM acetosyringone), were inoculated using a 25:1 dilution of the overnight culture (50 mL total). The larger culture was incubated at 30 °C, 225 rpm, overnight. Cells were harvested by centrifugation at 3,200g for 10 minutes. Cell pellets were resuspended in 1 volume of 10 mM MES, pH 5.6, 10 mM MgCl_2_. Cells were gently vortexed to resuspend the pellet. Cell suspensions were centrifuged at 3,200g for 10 minutes. Cell pellets were resuspended in 10 mL of 10 mM MES, pH 5.6, 10 mM MgCl_2_. The OD600 values were measured for each of the cultures. Cell suspensions were diluted using the same buffer to an OD600 of 1. Acetosyringone was added to the pTRV1 cell suspension to a final concentration of 400 µM. The different pTRV2-LIC constructs were mixed into 50 mL conical tubes with an equal volume of pTRV1 suspension, resulting in a final acetosyringone concentration of 200 µM.

Individual seedlings were inoculated through the abaxial side of the cotyledon. Plants were incubated at 24 °C and covered to avoid light for 24 hours and returned to 16/8h day-night cycles at the same temperature. Approximately 3 weeks later, the 4^th^ true leaf of each plant was sampled for acylsugars and RNA using a bisected leaf for each experiment. We found that inoculation timing is very important; the cotyledons should be inoculated after they have expanded, but before the first two true leaves have fully emerged.

Sample leaves of VIGS plants were dried for 1 week and weighed. Because the acylsugars in the samples were very concentrated, risking MS detector saturation, the acylsugar +2C isotope peak area was quantified by Quanlynx and normalized by internal standard and leaf weight. Ratios of tri- to tetraacylinositols were compared between TAIAT – P2/P1 and empty vector plants. Welch’s t-test was used to compare the tri- to tetraacylinositol ratio of TAIAT-targeted and empty vector plants. Raw data are available in Supplemental Table S2.

### qPCR analysis

RNA was extracted with the RNeasy Plant Mini Kit including on-column DNase digestion (Qiagen, Hilden, Germany), according to the manufacturer’s instructions. RNA was quantified with a Nanodrop 2000c instrument (Thermo Fisher Scientific, Waltham, MA). cDNA was synthesized using 1 µg of the isolated RNA and SuperScript II Reverse Transcriptase (Invitrogen, Carlsbad, CA). The cDNA samples were diluted 200-fold (10-fold initial dilution and 20-fold dilution into qPCR reactions). qPCR reactions (10 µL) were created with SYBR Green PCR Master Mix (Thermo Fisher Scientific, Waltham, MA) and primers were used at a final concentration of 200 nM. RT_TAIAT_F/R, RT_Actin_F/R, and RT_EF1α_F/R primers were used to detect *TAIAT, Actin*, and *EF1α* transcripts, respectively (Supplemental Table S3). Reactions were carried out with a QuantStudio 7 Flex Real-Time PCR System (Applied Biosystems, Warrington, UK) by the Michigan State University RTSF Genomics Core. The following temperature cycling conditions were applied: 50°C for 2 minutes, 95°C for 10 minutes, 40 cycles of 95°C for 15 seconds and 60°C for 1 minute. Relative expression of *TAIAT* was calculated with the ΔΔCt method (Pfaffl, 2001), and normalized to the geometric mean of *Actin* and *EF1α* transcript levels. The mean expression values of the transcripts in the control plants were used for normalization. 3-4 technical replicates were used for all the qPCR reactions.

### Purification of *myo*-inositol acylsugars and analysis by NMR spectroscopy

For metabolite profiling and purification, aerial tissues of 20 ten-week-old *S. quitoense* plants (∼0.25 m height) were extracted in 1.9 L of acetonitrile: isopropanol (AcN:IPA, v/v, 1:1) for 10 mins (plants were cut at the stems and stem junctions). Approximately 1 L of the *S. quitoense* bulk extract was concentrated to dryness under vacuum, redissolved in 5 mL of AcN:IPA, and fractionated by repeated injection of 200-µL aliquots onto a Thermo Scientific Acclaim 120 C18 HPLC column (4.6 x 150 mm, 5 µm particles) with reversed-phase gradient elution (0.15% formic acid and acetonitrile) and automated fraction collection. HPLC fractions of sufficient purity of a single metabolite, as assessed by LC/MS, were combined and concentrated to dryness. Samples were dissolved in acetonitrile-*d*_3_ (99.96 atom % D) and transferred to solvent-matched Shigemi tubes or Kontes tubes for NMR analysis. 1D (^1^H, ^13^C and TOCSY) and 2D (gCOSY, gHSQC, gHMBC, *J*-resolved, TOCSY and ROESY) NMR spectroscopic techniques served as the basis for structure elucidation of purified *S. quitoense* acylsugars. ^1^H, ^13^C, gCOSY, gHSQC, coupled-gHSQC, gHMBC, *J*-resolved, TOCSY and ROESY NMR experiments were performed using a Bruker Avance 900 MHz spectrometer equipped with a TCI triple resonance probe or an Agilent DDR2 500 MHz spectrometer equipped with OneNMR probe (with Protune accessory for hands-off tuning). 1D-TOCSY transfer experiments were performed using a Varian Inova 600 MHz spectrometer equipped with a Nalorac 5 mm PFG switchable probe (pretuned for ^1^H and ^13^C). All spectra were referenced to non-deuterated solvent signals: acetonitrile-*d*_3_ (^δ^H = 1.94 and ^δ^C = 118.70 ppm). NMR spectra were processed using TopSpin 3.5pl7 or MestReNova 12.0.0 software. Because these metabolites were identified without authentic standards or synthetic confirmation, their structures should be considered putative, meeting level 2 criteria of the Metabolomics Standards Initiative guidelines (Sumner et al., 2007). Further evidence for structural assignments described herein are outlined in (Hurney, 2018). The basis for the clockwise carbon numbering system used for assignment of *myo*-inositols in this study traces from the biosynthetic conversion of D-glucose 6-phosphate to 1L-*myo*-inositol-1-phosphate (Loewus and Murthy, 2000). Due to the plane of symmetry of the *myo*-inositol ring system about the C2 and C5 positions, an enantiomeric assignment is possible by substituting the C1 and C3 positions (1D-*myo*-inositol), followed by counterclockwise numbering. Due to the scarcity of purified material (most < 3 mg), absolute stereochemical configurations were not determined.

## Accession numbers

ASAT4 homolog sequence data is available in Genbank: TAIAT, (MT024677); c38687_g1_i1, (MT024678); c38687_g2_i1, (MT024679).

## Acknowledgements

We thank staff of the MSU RTSF Mass Spectrometry and Metabolomics facility for analytical chemistry guidance. We also are grateful to Eran Pichersky and Kevin Walker for helpful advice.

## Supplemental Data

**Supplemental figure S1. Negative-ion MS-MS spectra of product ions generated from [M+formate]**^**-**^ **of *S. quitoense* acylinositols reveal C10 and C12 fatty acid carboxylate ions.**

**Supplemental figure S2. Positive-ion MS-MS spectra of product ions generated from [M+NH**_**4**_]^**+**^ **of *S. quitoense* acylinositols.**

**Supplemental figure S3. Multiple sequence alignment of sequences used in phylogenetic analysis.**

**Supplemental figure S4. qRT-PCR analysis of TAIAT transcript abundance in VIGS plants.**

**Supplemental figure S5. ESI+ mode non-mass selective fragmentation of forward enzyme assay products of TAIAT (I4:24 (2**,**2**,**10**,**10) and I4:26 (2**,**2**,**10**,**12)) generated from incubations of triacylinositols I3:22 and I3:24 with acetyl CoA.**

**Supplemental figure S6. Co-elution analysis of *in vitro* assay products of TAIAT using C18 and F5 chromatography.**

**Supplemental figure S7. *In vitro* assays comparing different ASAT4 homolog activities with I3:22 (2**,**10**,**10).**

**Supplemental figure S8. *In vitro* assays comparing different ASAT4 homolog activities with I3:24 (2**,**10**,**12).**

**Supplemental figure S9. *In vitro* reverse reactions of TAIAT.**

**Supplemental figure S10. Positive-ion mode mass spectra with ramped collision voltage (10-60 V) of triacylinositol products generated from reverse enzyme assay products of TAIAT with I4:24 and I4:26 (products are I3:22 (2**,**10**,**10) and I3:24 (2**,**10**,**12)).**

**Supplemental Table S1. NMR chemical shifts for four acylinositols from S. quitoense.**

**Supplemental Table S2. Acylsugar analysis of S. quitoense VIGS plant samples.**

**Supplemental Table S3. Oligonucleotides used in this study.**

